# Minicollagens play the governing role in polar capsule formation in parasitic cnidarians, the Myxozoa

**DOI:** 10.1101/2022.03.16.484579

**Authors:** Jiří Kyslík, Marie Vancová, Pavla Bartošová-Sojková, Alena Lövy, Astrid S. Holzer, Ivan Fiala

## Abstract

Minicollagens are major structural components in the biogenesis of nematocysts in Cnidaria. Sequence mining and recent proteomic analysis of polar capsules, homologues of cnidarian nematocysts, have confirmed the presence of minicollagens in this evolutionarily ancient cnidarian endoparasitic group. Nonetheless, the role of nematocyst-associated proteins in polar capsule morphogenesis has never been studied in myxozoans. Here, we report the gene expression of three myxozoan minicollagens, *ncol-1*, *ncol-3*, and the recently identified *ncol-5*, during the intrapiscine development of *Myxidium lieberkuehni*, the myxozoan parasite of Northern pike *Esox lucius*. Moreover, we determined the abundance and localisation of Ncol-1 and Ncol-5 proteins in the developing myxozoan stages by western blotting and by immunofluorescence and immunogold electron microscopy. We found that expression of minicollagens was spatiotemporally restricted to developing polar capsules in sporogonic stages. Intriguingly, Ncol-1 and Ncol-5 were localised as major components of the polar capsule wall and polar tubule. These results support the common origin of nematocysts and myxozoan polar capsules. Furthermore, our findings have practical implications for a more accurate identification of developmental stages of myxozoan parasites.

## Introduction

Nematocysts, a clear synapomorphy of the phylum Cnidaria, represent a single-celled innovation used for defensive and predatory strategies (Mariscal, 1974; Özbek et al., 2009). The weapon-like fashion of discharge of these organelles is among the fastest known in any living organism (Holstein and Tardent, 1984; Nuchter et al., 2006). Nematocysts have evolved extensive phenotypic and functional complexity during evolution (David et al., 2008). Due to their large diversity, nematocysts also serve as important taxonomic markers in the classification of cnidarians (Weill, 1934; Mariscal, 1974). The morphogenesis of nematocytes originates from the proliferation of the neuronal stem cell population (Tardent, 1995). After that, the nematocytes form the primordial nematocyst capsule in the post-Golgi vacuole via vesicle fusion (Holstein, 1981). The process of nematocyst maturation is preceded by continuous formation of the capsule wall, followed by tubule formation and final invagination into the capsule body (Engel et al., 2001; Adamczyk et al., 2010).

Myxozoa, a parasitic group of cnidarians, produce spores with simplified nematocysts called polar capsules that play an important role in the attachment of the parasite to the host (El-Matbouli et al., 1999; reviewed in Americus et al. 2020). Although free-living cnidarians and parasitic myxozoans possess distinct body forms, striking morphological similarities of nematocysts and polar capsules suggested that Myxozoa are Cnidaria (Lom and de Puytorac, 1965), which was later confirmed by molecular analysis of nematocyst-related genes (Holland et al., 2011). Myxozoa represent an evolutionarily ancient group of parasites that mostly invade fish and annelid hosts and that originated around ~651 Mya, after the emergence of the main cnidarian lineages (Holzer et al., 2018). They underwent dramatic genome and cellular reduction (Alama-Bermejo and Holzer, 2021). Prior to the development of myxospores with polar capsules, a unique cell-in-cell formation and series of presporogonic stages are produced. Multicellular proliferative stages, plasmodia, emerge at the final site of infection (Lom and Dyková, 2006). The plasmodium undergoes intense cell differentiation to form sporogonic cell stages destined for the final spore production. Before polar capsule formation, capsulogenic cells represent cellular precursors for the formation of polar capsules (Lom and Dyková, 2006). As in more complex nematocysts, polar capsule development precedes the simultaneous formation of the external polar tubule and capsular primordium with final tubule invagination (Desser et al., 1983; Casal et al., 2002).

By studying the molecular components of nematocysts, a number of nematocyst-specific structural proteins, such as minicollagens, nematogalectins, and NOWA, have been identified (Engel et al., 2001; Engel et al., 2002; Özbek et al., 2004; Hwang et al., 2010; Adamczyk et al., 2008; Adamczyk et al., 2010). Minicollagens are important components of the nematocyst with a typical protein domain architecture consisting of a truncated central collagen domain flanked by poly-proline stretches of varying sizes that terminate with cysteine-rich domains (CRD) (Özbek et al., 2002). The expression pattern and subcellular localisation revealed that minicollagens are present only in developing nematocytes, whereas they cannot be detected with the respective antibodies in mature nematocysts (Engel et al., 2001; Adamczyk et al., 2008). In *Hydra*, the highly conserved minicollagen 1 (Ncol-1) is restricted to the capsule wall, whereas Ncol-15 is present in the nematocyst tubule (Engel et al., 2001; Adamczyk et al., 2008). Nematocyst capsule assembly undergoes a disulfide-dependent process of CRD integration of minicollagens with variable intermolecular cross-linking and polymerisation kinetics (Tursch et al., 2016).

In contrast to the molecular complexity of regular cnidarian nematocysts (Rachamim et al., 2015) the myxozoan polar capsule gene repertoire shows dramatic reduction (Shpirer et al., 2014; Piriatinskiy et al., 2017). However, a recent study describing a new type suggested the minicollagen repertoire akin to cnidarians (Kyslík et al., 2021). The identification of minicollagen 5 (Ncol-5) revealed a unique noncanonical domain architecture with remarkable substitutions of poly-proline stretches to glycine/serine-rich derived domain patterns. A unique function of Ncol-5 in polar capsule or development of polar tubule has been suggested (Kyslík et al., 2021), but the exact structural role of minicollagens in myxospore development has so far been not investigated.

Here, we report the gene expression profile and localisation of minicollagens during polar capsule development in *Myxidium lieberkuehni* Bütschli, 1882, a myxozoan parasite of Northern pike *Esox lucius* L. To understand the distribution of minicollagens in polar capsule morphogenesis, we raised polyclonal antibodies against Ncol-1 and the recently identified Ncol-5 and localised both proteins in parasite cells during its intrapiscine development. Both minicollagens showed stage-specific gene expression exclusively related to the spore-forming stages and both proteins were located in the polar capsule wall and tubule within developing polar capsules, as observed for cnidarian nematocysts (Engel et al., 2001; Adamczyk et al., 2008). Interestingly, antibodies generated against Ncol-1 and Ncol-5 recognised the same structures within the polar capsule, despite their differing domain architecture. Even though the morphology of myxozoan parasites is significantly reduced, polar capsules seem to have retained a similar architecture and molecular structure as nematocysts of free-living cnidarians.

## Results

### Minicollagens are expressed exclusively in myxozoan spore-forming stages

Regular screening of the urinary bladder of pikes revealed the presence of small presporogonic stages predominant in the host, in August and September (Fig. 1A-B), which had grown into large plasmodial stages in October (Fig. 1C-D) with budding of daughter plasmodia (Fig. 1D). Starting in January, we detected the formation of pansporoblasts with developing myxospores in large plasmodia (Fig. 1E); mature spores began to appear freely floating in the urinary bladder in April (Fig. 1F-H).

**Fig.1.**
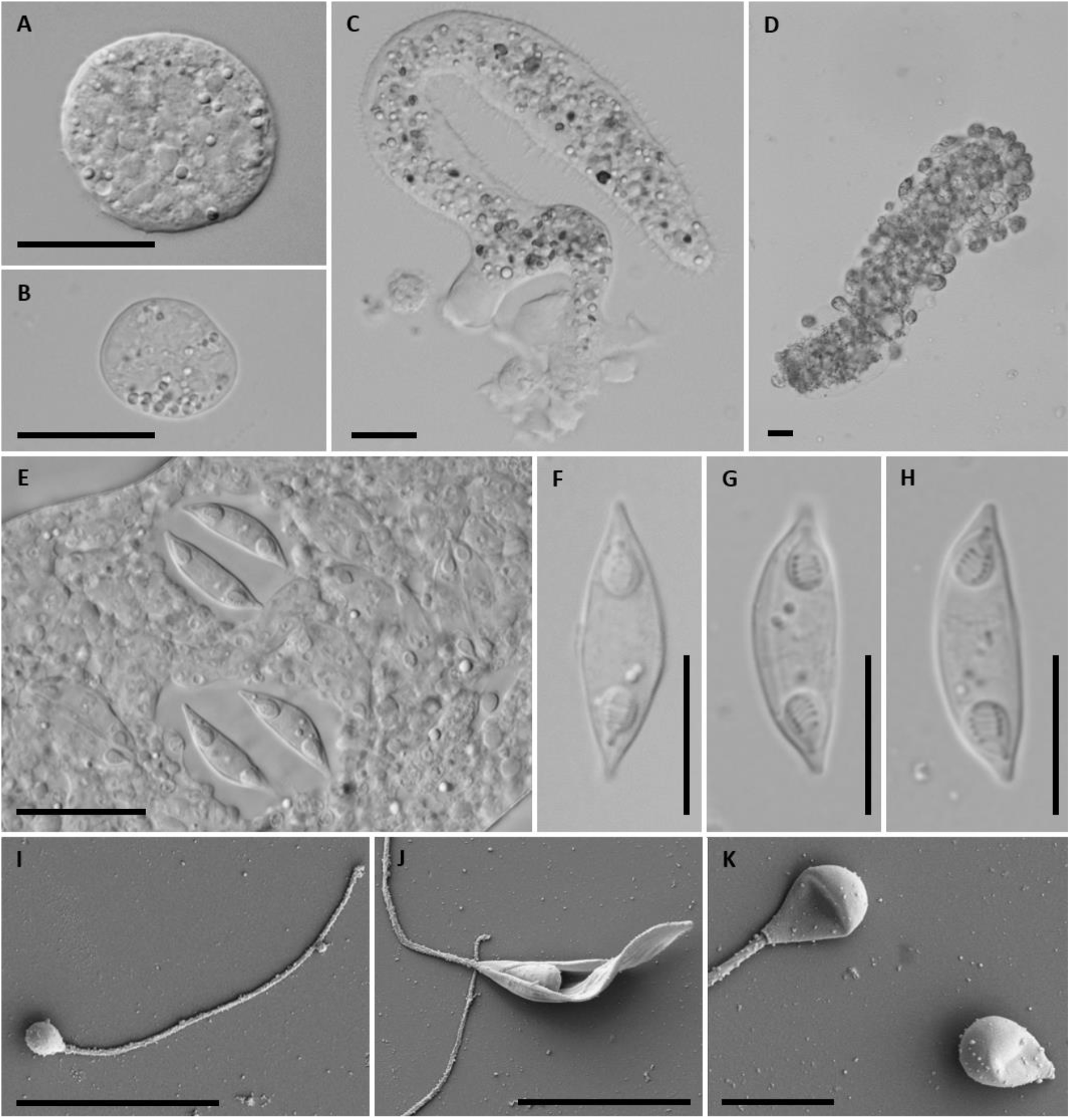
Light microscopy images of developmental stages of *Myxidium lieberkuehni*. **(A–B)** Early plasmodial stages. **(C–D)** Growing plasmodia with budding of daughter plasmodia in (D). **(E)** Pansporoblasts with two mature myxospores in each of them. **(F–H)** Mature myxospores. **(I–K)** Scanning electron microscopy of polar capsules with discharged polar filaments and a visible spore valve (J). (Scale bars: A–E: 20 μm; F–K: 10 μm).

All three minicollagens were exclusively expressed in spore-forming stages collected between January and April, while the gene expression in other months (August to December) was negligible (Fig. 2A–C). The highest level of relative expression of all three minicollagens was on average 1000 times higher in sporogonic stages compared to presporogonic stages (August to December) (Fig. 2D). The peak expression level of all minicollagens was detected in March, during pansporoblast formation typical for myxospore development (Fig. 2A–C). In the developmental stages with mature myxospores (April), a decrease in minicollagen expression (1–2.5 times for *ncol-1*; 1.2–72 times for *ncol-3*; 1.2–12 times for *ncol-5*) was detected compared to the sporogonic phase in the previous two months (February and March) (Fig. 2A–C). Overall, all minicollagens showed a co-linear expression during the myxospore development (Fig. 2E).

**Fig.2.**
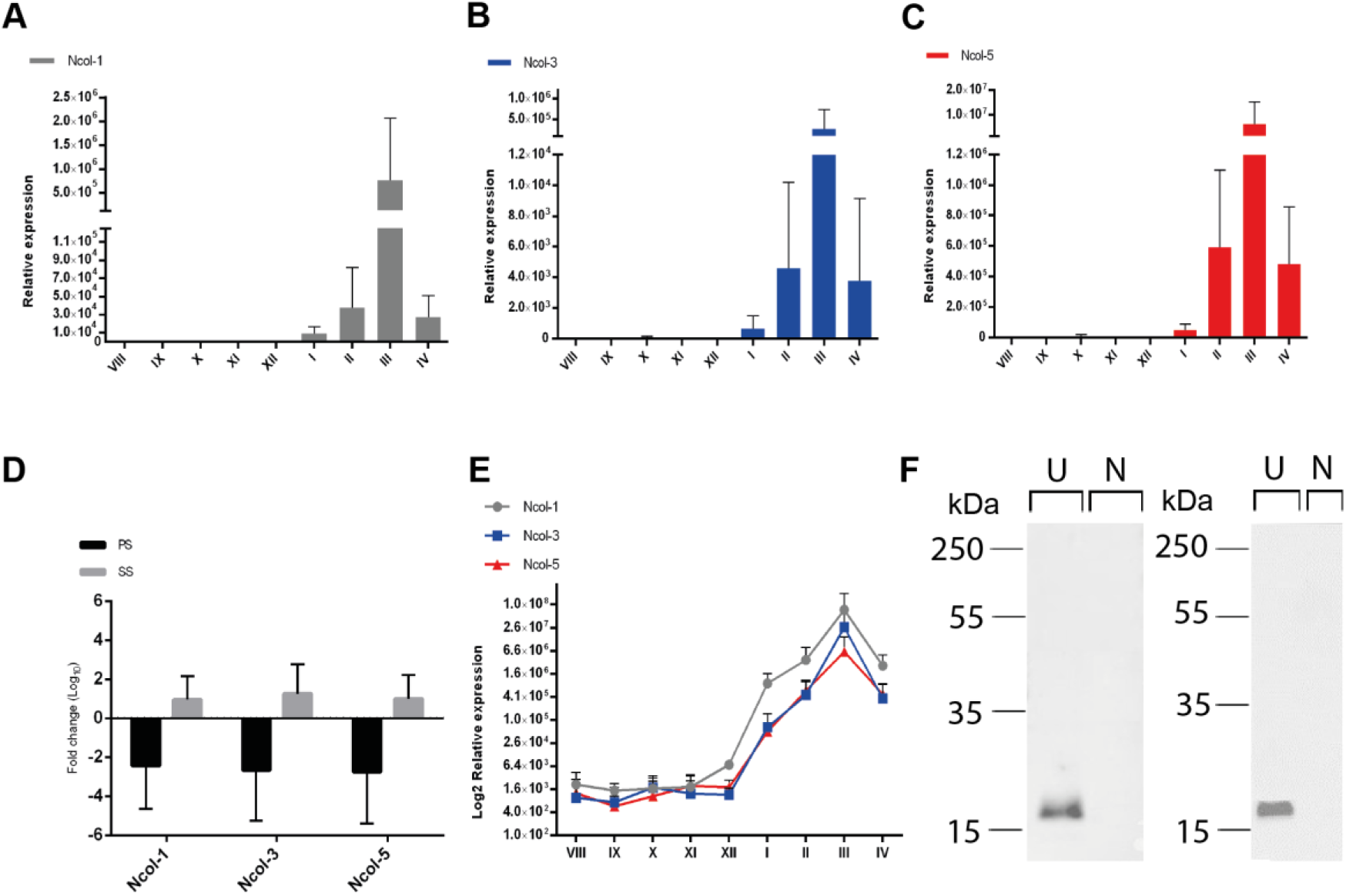
Gene expression profiling of *Myxidium lieberkuehni* minicollagens (*ncol-1*, *ncol-3*, *ncol-5*) during the myxospore development. **(A–C)** Gene expression of *ncol-1* (grey), *ncol-3* (blue), and *ncol-5* (red) relative to two housekeeping genes during myxospore development. Broken X-axis: month of collection, Y-axis: relative expression values. **(D)** Fold change expression (log_10_) of minicollagens in presporogonic stages (PS, black bars) and sporogonic stages (SS, grey bars). **(E)** Collinear expression of *ncol-1* (grey), *ncol-3* (blue), and *ncol-5* (red) relative to two housekeeping genes (log_10_) during myxospore development. X-axis: month of collection, Y-axis: log_10_ transformed relative expression values. **(F)** Western blots of rabbit polyclonal anti-Ncol-1 (left panel) and anti-Ncol-5 (right panel) in lysates of sporogonic stages (U) and control host tissue (N), each from one fish individual.

### Minicollagens are abundant proteins in myxozoan sporogonic stages

The efficiency and specificity of the custom antibodies was evaluated by western blotting using parasite lysates prepared from *M. lieberkuehni* developmental stages containing polar capsules. A major band with a molecular weight of ~17 kDa was detected against either anti-Ncol-1 or anti-Ncol-5 antibody (Fig. 2F). We detected a shift in the molecular weight of the actual protein in comparison to the predicted protein size - 14 kDa for both proteins. Both antibodies showed a high specificity towards parasite minicollagens with no cross-reactivity detected between anti-Ncol-1 and anti-Ncol-5 or with the host muscle tissue lysates (Fig. 2F).

Strong minicollagen signal was detected only in sporogonic stages (Fig. 2F) while no reaction was observed in presporogonic stages (data not shown).

### Minicollagens exhibit post-translational modifications

Based on the difference between the predicted and actual protein sizes, post-translational modifications (PTMs) of primary amino acid sequences of both proteins were predicted. Ncol-1 showed one O-glycosylation site (Thr44) in the N-terminal CRD and two O-glycosylation sites (Thr156; Ser160) in the C-terminal CRD (Fig. S2A). Considering the presence of poly-proline repeats in both N/C-terminal parts and in a central collagen domain (Gly-X-Y), a substantial number of hydroxylation sites (37) in Ncol-1 were predicted (Fig. S2A). Ncol-5 exhibited four O-glycosylation sites (Ser48; Ser52; Ser 56; Ser60) in the N-terminal glycine-serine rich domain (Fig. S2B), a region that is highly divergent from other myxozoan minicollagens, including Ncol-1. Furthermore, several hydroxylation sites (19) in proline residues within the N/C terminal part, including a collagen domain (Gly-X-Y), were predicted for Ncol-5. N-linked glycosylations were not detected.

### Ncol-1 and Ncol-5 are both located in the wall and tubule of the polar capsule

Morphogenesis of *M*. *lieberkuehni* polar capsules was examined using labelling with antibodies directed towards Ncol-1 and Ncol-5 minicollagens and visualised by confocal laser scanning microscopy. Both antibodies recognised the proteins within the developing polar capsules in maturing spores inside pansporoblasts (Fig. 3A & Fig. 5A). They both recognized the proteins in the wall and polar tubule of the capsule throughout capsule formation (Fig. 3B & Fig. 5B). Minicollagens were proven to be present in the developing primordial polar capsules during myxospore formation, whereas the proteins in mature myxospores were not labelled (Fig. 3A & Fig. 5A; shown by arrowhead). No Ncol-1 and Ncol-5 signal was observed in presporogonic stages (data not shown), which corresponds to the lack of gene expression in these stages (Fig. 2A-C & Fig. 2D, E). Autofluorescence of crystal-like structures (Fig. S3A) was observed in the presporogonic stages under epifluorescence (Fig. S3B).

**Fig. 3.**
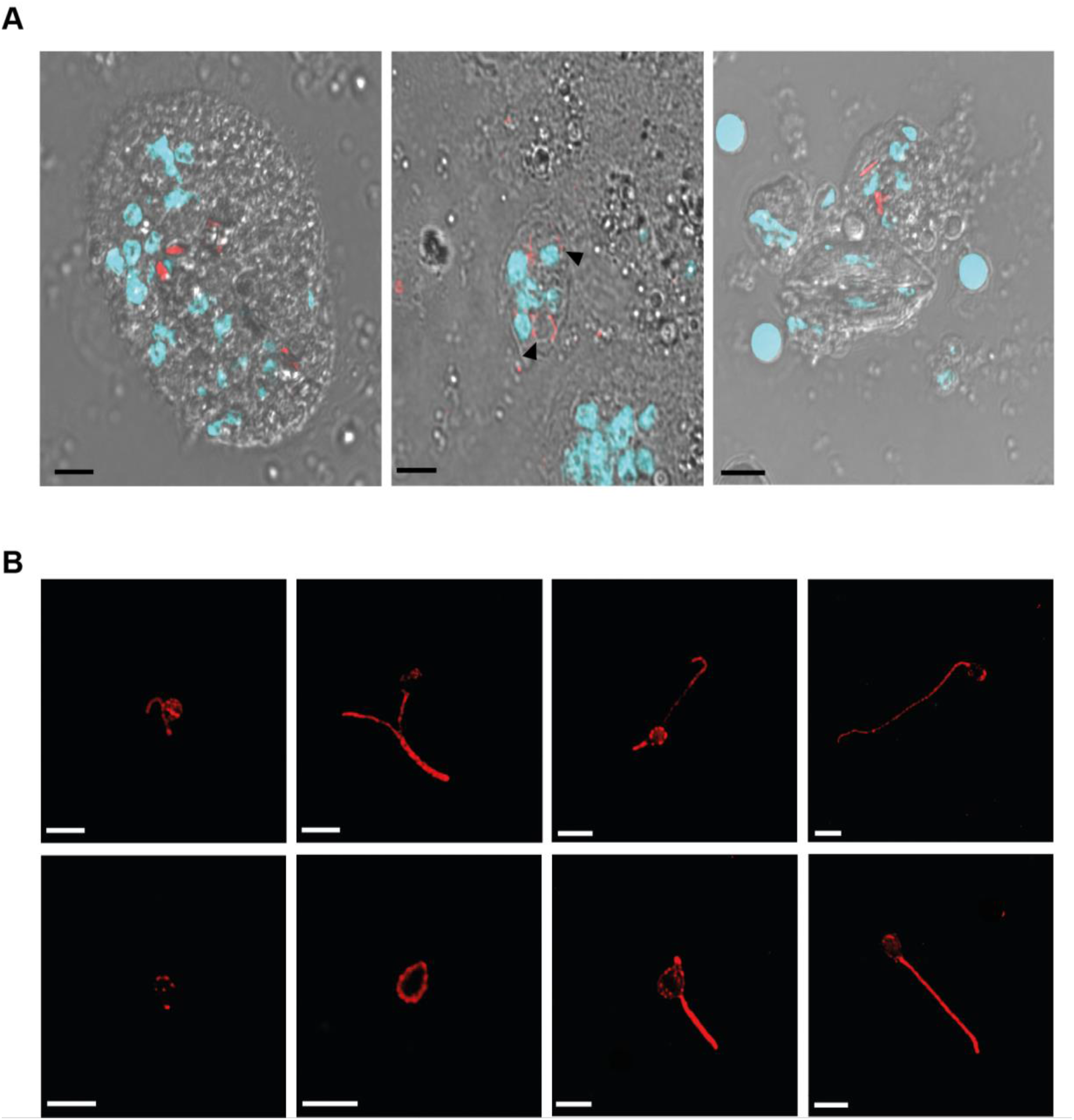
Immunolabelling of Ncol-1 in *Myxidium lieberkuehni* sporogonic stages and in developing polar capsules. **(A)** Confocal images of sporogonic stages: from left to right developing pansporoblast, immature polar capsules (arrowheads) within the developing spores, the late stage with mature polar capsules in which DNA (DAPI; light blue) and Ncol-1 (Alexa Fluor Plus™ 647; red with arrows) are visualized and merged with the differential interference contrast image; **(B)** confocal images of Ncol-1 in extracted polar capsules in which Ncol-1 (Alexa Fluor Plus™ 647; red) is visualized (Scale bar: 5 μm).

Detailed localisation of minicollagens during polar capsule formation was demonstrated by immunogold electron microscopy (EM). Ncol-1 and Ncol-5 antibodies localised both minicollagens in the wall and tubule of the polar capsules (Fig. 4 & Fig. 6). During myxospore formation, antibodies revealed the accumulation of signals in the matrix of the capsular primordium (Fig. 4A–C & Fig. 6C) and a gradual condensation either in the polar capsule wall or in the polar tubule (Fig. 4B–E & Fig. 6A–B, D, E). The polar tubule formation was never observed, except for a coiled polar filament (Fig. 4D & Fig. 6E) that was already present in the final structure of the polar capsule body, and both minicollagens were located in the tubule structure (Fig. 4D, E & Fig. 6E). No signal against the minicollagens was detected outside the polar capsule in the parasite myxospores as well as in the spore walls.

**Fig. 4.**
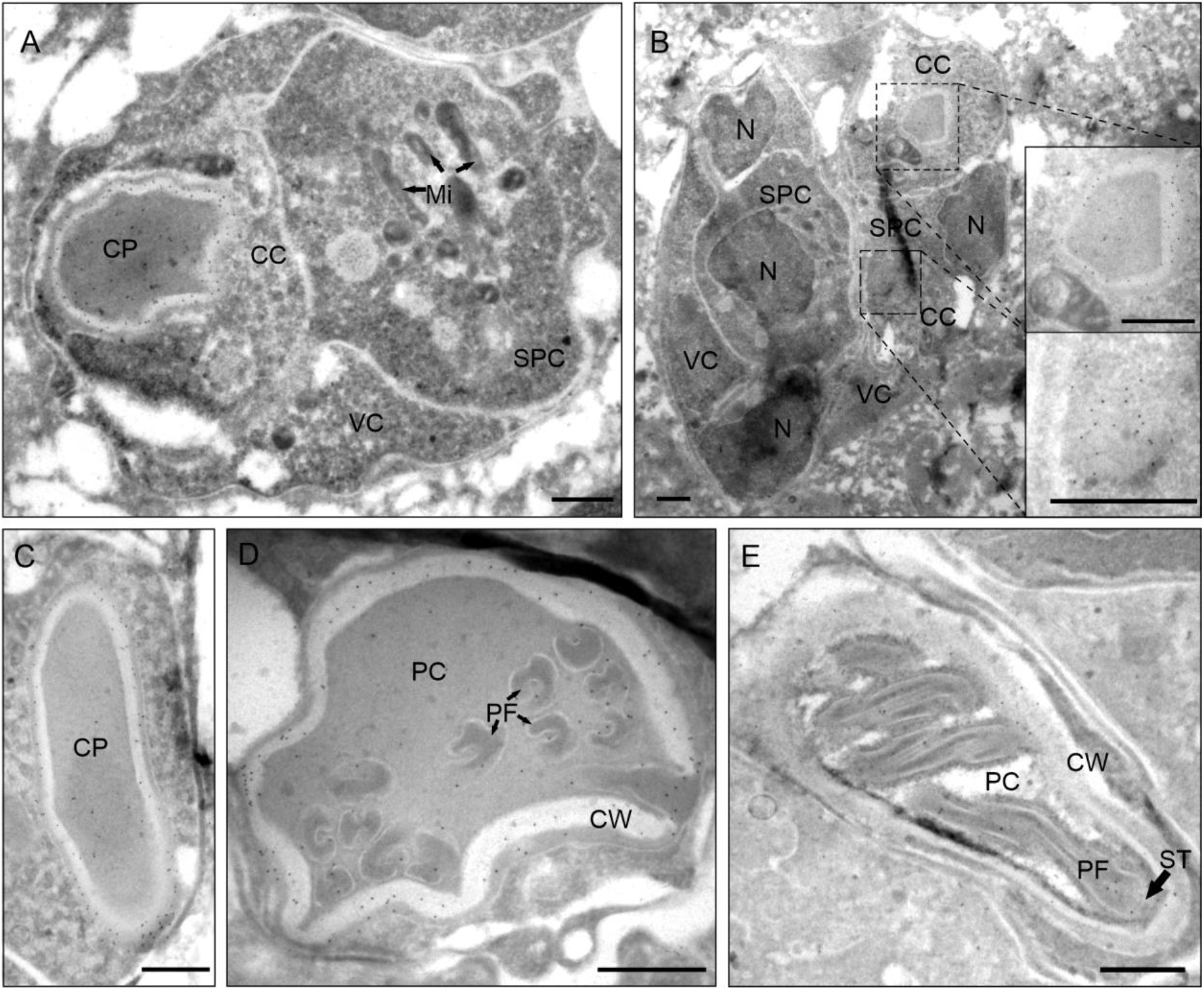
Transmission electron microscopy (TEM) of immunogold-labelled Ncol-1 during the *Myxidium lieberkuehni* polar capsule development. Immunogold-labelling of Ncol-1 (indicated by black dots) in: **(A–B)** Pansporoblasts containing the capsulogenic cell (CC) with the capsular primordium (CP), the sporoplasm cell (SPC), mitochondria (Mi, arrows), nuclei (N), and the valvogenic cell (VS). Enlarged images of developing capsular primordium with Ncol-1 signal during formation of the capsular wall (top) and matrix (bottom) in **(B)**. **(C)** Capsular primordium (CP) with the signal in the capsular matrix and the wall. **(D–E)** Polar capsule sections (PC) with the signal in both coiled polar filament (PF, arrows) and capsular wall (CW) with a stopper (ST, arrow). (Scale bar: 500 nm).

**Fig. 5.**
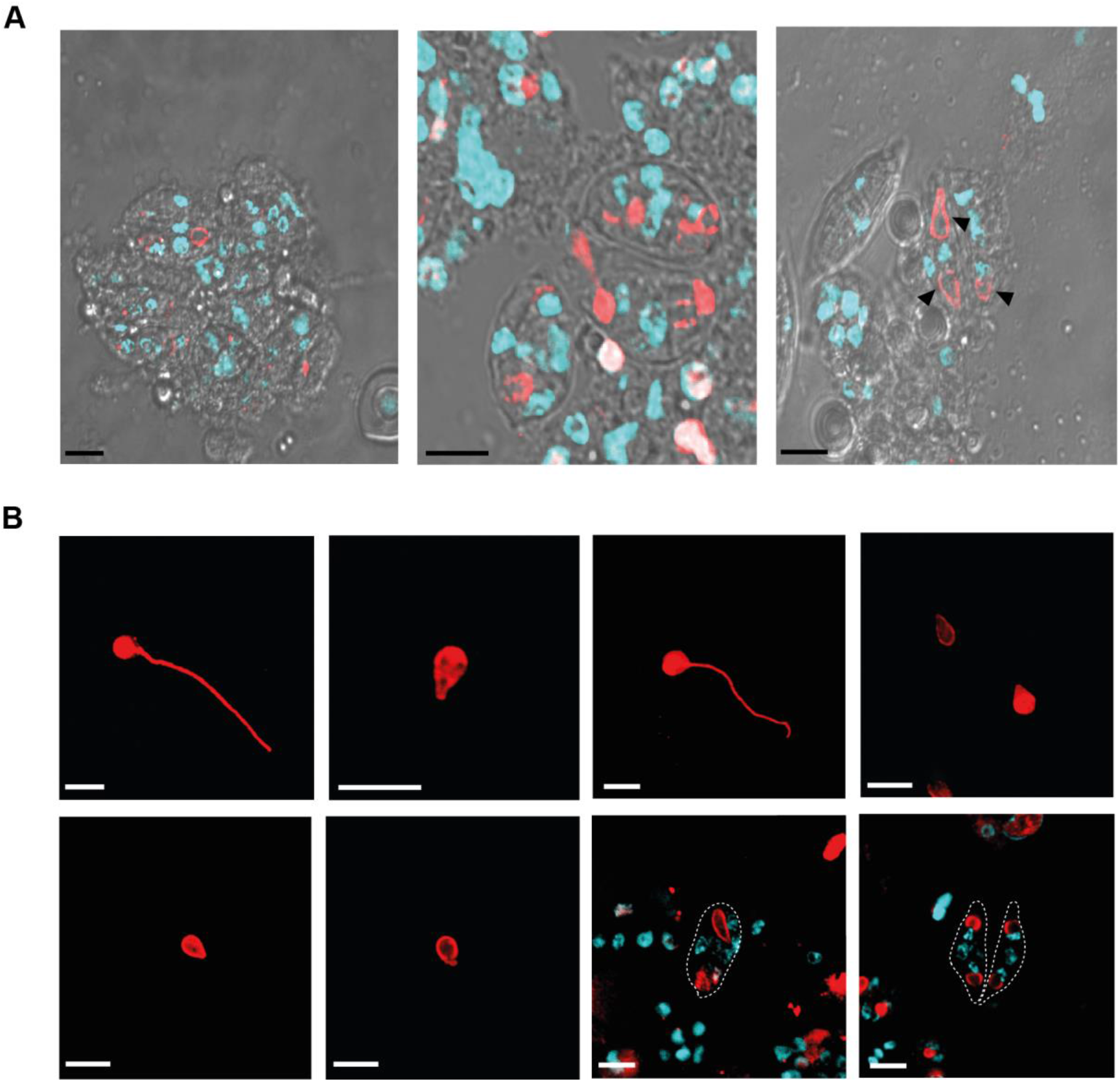
Immunolabelling of Ncol-5 in *Myxidium lieberkuehni* sporogonic stages and in polar capsules. **(A)** Confocal images of sporogonic stages: from left to right differentiating pansporoblasts, early stages of developing spores with immature polar capsules, late stage with developing polar capsules (arrowheads) in which DNA (DAPI; light blue) and Ncol-5 (Alexa Fluor Plus™ 647; red) are visualized and merged with the differential interference contrast image. **(B)** Confocal images of extracted polar capsules in which DNA (DAPI; light blue) and Ncol-5 (Alexa Fluor Plus™ 647; red) are visualized and merged. (Scale bar: 5 μm).

**Fig. 6.**
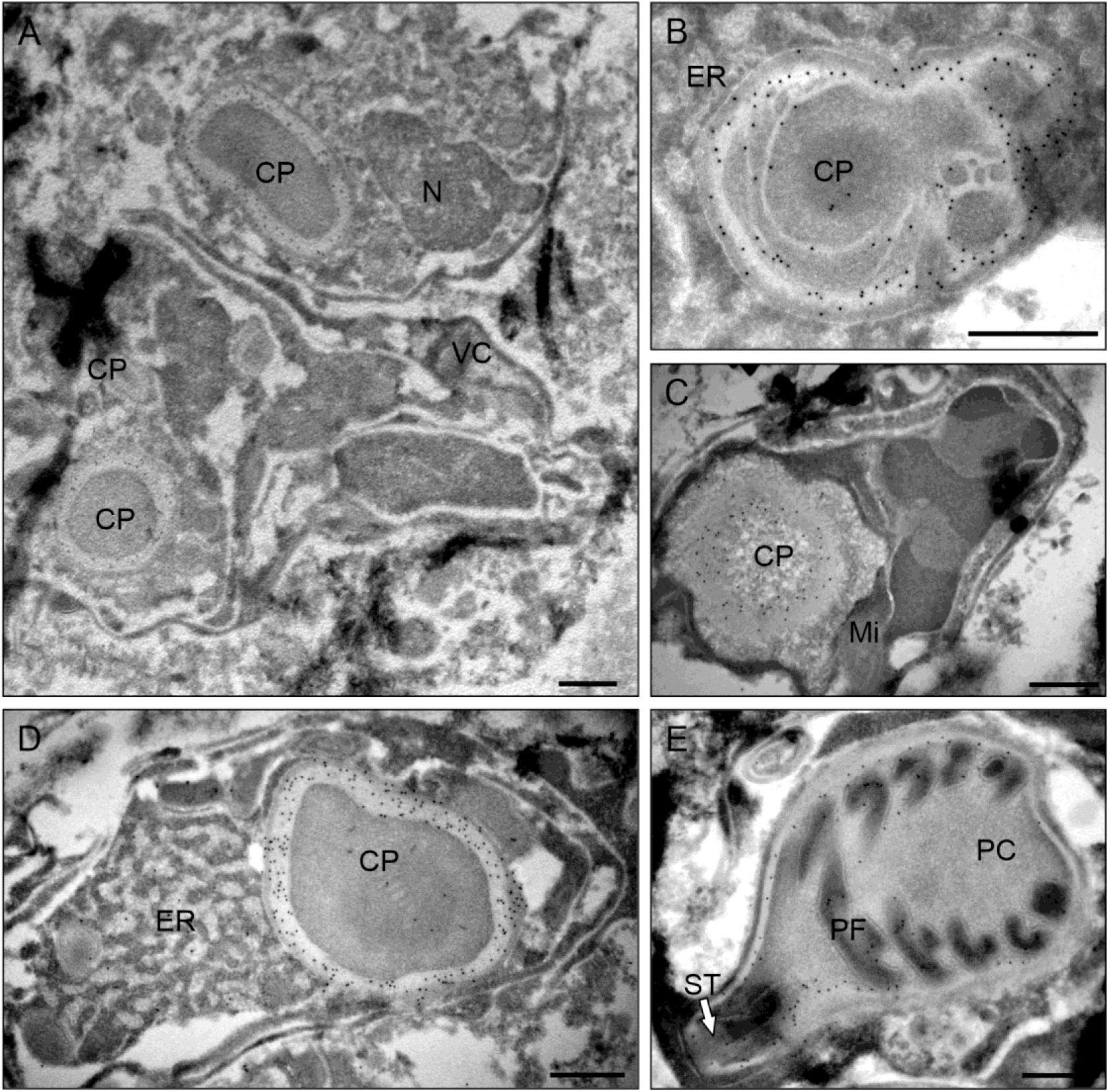
Transmission electron microscopy (TEM) of immunogold-labelled Ncol-5 during the *Myxidium lieberkuehni* polar capsule development. Immunogold-labelling of Ncol-5 (indicated by black dots). **(A)** Pansporoblast containing the capsulogenic cell with a capsular primordium (CP), the nucleus (N), and the valvogenic cell (VS). **(B)** Ncol-5 signal in developing wall of the capsular primordium (CP) surrounded by endoplasmic reticulum (ER). **(C)** Capsular primordium (CP) within the polar capsule with the signal in the capsular matrix and the wall. **(D–E)** Polar capsule sections (PC) with the signal in both coiled polar filament (PF) in (E), capsular wall (CW) and primordium (CP) with a stopper (ST, arrow). (Scale bar: 500 nm).

## Discussion

The role of minicollagens in the morphogenesis of cnidarian nematocysts has been investigated in free-living cnidarians in several studies (Engel et al., 2001; Adamczyk et al., 2008; Zenkert et al., 2011; Beckmann and Özbek, 2012; Tursch et al., 2016). However, localisation of minicollagens within the myxozoan organelle, the polar capsule, has not yet been characterised. For the first time in myxozoans, we report the kinetics of minicollagen expression of all three minicollagens identified in *Myxidium lieberkuehni* and investigate the localisation of two of them (Ncol-1 and Ncol-5) in polar capsule development. Myxozoan Ncol-5, a new type of minicollagen, was identified with a unique gene structure that suggested a specific function in myxospore development (Kyslík et al., 2021).

Alongside determining the expression and localisation of minicollagens in polar capsule morphogenesis, we scrutinized the previously underexplored development of *M. lieberkuehni* in the fish host, terminated by the production of myxospores (Debaisieux, 1920; Bremer, 1922). The morphogenesis observed in *M. lieberkuehni* is consistent with the general myxosporean development described in previous studies (e.g. Weill, 1934; Desser et al., 1983; Casal et al., 2002, Okamura et al. 2015, Morris et al. 2010). We detected the initial pansporoblast formation in January, which continues until April when many mature spores containing polar capsules are discernible. During this season, remarkably high expression levels of the minicollagens (*ncol-1*, *ncol-3*, *ncol-5*) were observed in sporogonic stages, due to the participation of studied genes in the formation of the polar capsules in developing spores, as generally reported from immature nematocysts in free-living cnidarians (Engel et al., 2001; Adamczyk et al., 2008; Hwang et al., 2010). In our myxozoan model organism, minicollagen expression began in January, when we first detected the formation of pansporoblasts in the plasmodia of *M. lieberkuehni*. The highest peak of gene expression of all assayed minicollagens was in March, supported by light microscopy observations showing the highest number of pansporoblasts and mature spores. In the following month, expression of minicollagens was downregulated along with lower pansporoblast formation. We were unable to provide data from May to July because fishing of the pike during the spawning season is unlawful. However, we assume that spore maturation terminates in May with an expected drop of minicollagen expression in this season. The life cycle of myxospores continues with the release of spores and by the development in a definitive, yet undiscovered, host of *M. lieberkuehni*, which is probably an oligochaete. The development of myxosporeans in their definitive hosts ends up with the formation of actinospores which also bear polar capsules (Wolf and Markiw, 1984). Thus, we assume a biphasic expression of minicollagens in the myxozoan life cycle. From our observations, we conclude that the actinosporean phase of *M. lieberkuehni* takes place from May to July at the latest, as we have already detected the development of early plasmodia in the urinary bladder of the pikes in August. Hence, we suppose a short-term expression of minicollagens occurs during an actinospore maturation, that probably takes place within several weeks compared to the development in the fish host (El-Matbouli and Hoffmann, 1998; Morris and Freeman, 2010).

Our results showed collinear expression of all three myxozoan minicollagens studied. Khalturin et al. (2019) found that minicollagen genes in representatives of major cnidarian lineages are organized into clusters with collinear expression. Our previous study of phylotypic clustering in myxozoan minicollagens identified juxtaposition of two minicollagens Ncol-1 and Ncol-4, while Ncol-2, Ncol-3, and Ncol-5 showed separated positions (Kyslík et al., 2021). Medusozoan genomes contain a greater number of minicollagen genes than those of the anthozoan *Nematostella vectensis*, suggesting a correlation with the higher complexity of the nematocyst repertoire in medusozoans compared to the anthozoans (Khalturin et al., 2019) and myxozoans, which have only one simple type. Therefore, a similar expression pattern of myxozoan minicollagens can be linked to the cell type-specific and spatiotemporal restricted expression during nematocyst development of the anthozoan *N. vectensis* (Babonis and Martindale, 2017; Sebé-Pedrós et al., 2018; Sunagar et al., 2018). Thus, we also predict some transcriptional regulation of myxozoan minicollagens during the myxospore development.

Ancient evolutionary origin and parasitism of Myxozoa (Holzer et al. 2018) might have led to the depletion of the nematocyst repertoire to a single type of polar capsule. Alternatively, one can suppose that continuous acquisition and complexity of novel nematocyst types (David et al. 2008) occurred after the split of free-living Cnidaria and Myxozoa. We assume that the parasitic lifestyle of early-branching Myxozoa did not require adaptations resulting in the diversification of nematocysts during the evolution. Notably, this is related to the utilization of myxozoan polar capsules exclusively for anchoring to the host contrary to prey capture and defence of cnidarian nematocysts. However, whether certain myxozoans possess additional types of polar capsule is questionable regarding the number of undescribed myxozoan species (Okamura et al., 2015). Although myxozoan polar capsules represent an adaptation of cnidarians to a parasitic life strategy (Piriatinskiy et al., 2017), our data support the conservation of protein function of minicollagens in myxozoans as observed in the nematocysts of free-living Cnidaria.

To verify the presence and abundance of minicollagen proteins during the parasite intrapiscine development, we first raised polyclonal antibodies against Ncol-1 and Ncol-5 and then examined them by western blotting. The minicollagens were detected in stages containing developing polar capsules while they were absent in presporogonic stages. Thus, in summary, minicollagen gene expression and protein production are unequivocally linked to *M. lieberkuehni* sporogenesis.

Despite the presporogonic stages being negative for minicollagens labelling, we noted internal crystal-like structures (Fig. S1A) showing autofluorescence (Fig. S1B). This signal might come from ingested host cells since plasmodia are an active feeding stage in Myxozoa (Current et al., 1979; Azevedo et al., 2013). We assume that observed crystals represent remnants of pigment granules, which include either lipofuscin or melanin contained in macrophages (Nakayama 2018).

Our finding of minicollagen expression and protein production during *M. lieberkuehni* sporogenesis corresponds to the presence of minicollagens in the proteome of polar capsules of another myxozoan *Ceratonova shasta* (Piriatinskiy et al., 2017). Both *M. lieberkuehni* minicollagens (Ncol-1, Ncol-5) showed a shifted band size (17 kDa) compared to their expected molecular weight (14 kDa) in Western blots (Fig. 1A). This suggested the presence of PMTs in studied proteins consistent with previous reports from cnidarians (Engel et al., 2001; Adamczyk et al., 2008; Gold et al., 2019). Our computational analyses of minicollagens performed to predict PMTs revealed that both proteins showed O-linked glycosylations and numerous hydroxylations (Fig. S2A, B), which may have caused larger actual protein sizes in Western blot assays in comparison to molecular weight predictions. Due to the insufficient number of polar capsule material, enzymatic deglycosylation/degradation assays need to be done to verify the PTMs predictions. Nevertheless, a substantial number of hydroxylations in either the central collagen domain (Gly-X-Y) or in poly-proline stretches might allow myxozoan minicollagens to form a triple-helical structure as reported in other cnidarians (Engel et al., 2001; Gorres and Raines, 2010).

Similar to gene expression and western blotting assays, immunolabelling at both light and TEM levels showed labelling of minicollagens in stages limited to sporogenesis. The inability of specific antibodies to bind the minicollagens in mature myxospores and their polar capsules could be caused by structural changes related to the final polymerisation of these proteins, as reported from cnidarian nematocysts (Engel et al., 2001; Engel et al., 2002; Özbek et al., 2002; Adamczyk et al., 2008) or by an insufficient permeability of hardened spore valves and associated problematic antibody entry into the myxospore itself. The latter is supported by the successful fluorescent visualization of minicollagens in polar capsules extracted from mature myxospores. Antibodies against Ncol-5 minicollagen recognized the same structures as Ncol-1, the wall and tubule of the polar capsules, in the fluorescent immunolabelling and immunogold EM. This co-distributional pattern indicates that Ncol-5, like other minicollagens, is involved in polar capsule formation, contrary to our previous suggestion of the unique, so far unknown, function of Ncol-5 exhibiting a noncanonical sequence structure (Kyslík et al., 2021). Immunogold staining of minicollagens also showed their accumulation of signal in the capsular matrix during the maturation of the polar capsule and signal condensation in the wall and polar tubule. This has also been observed in cnidarian minicollagens during the maturation of nematocysts (Engel et al., 2002). Although we clarify the localisation of minicollagens in developing polar capsules, the structural assembly of minicollagens during the development of polar capsules remains to be confirmed. Though previously reported from myxozoans (Westfall 1966; Desser et al., 1983), formation of external tubules was not detected for *M. lieberkuehni* in our ultrathin TEM sections, which could be due to the rapid tubule morphogenesis and invagination into the capsule, as shown in Cnidaria (Hwang et al., 2010), Based on the co-distribution of Ncol-1 and Ncol-5, a similar localisation of minicollagens can be assumed for other myxozoan species based on a common domain architecture of these genes, similar to that of Cnidaria (Zenkert et al., 2011).

Taken together, we describe the expression and localisation of minicollagens during myxozoan polar capsule development. Based on our findings, structurally different minicollagens (Ncol-1, Ncol-5) were recognised in the same locations with a co-distributional pattern in the polar capsule. Notably, both minicollagens are localised in the wall and tubule of the polar capsule. Our results represent a pioneering study elucidating the localisation of structural proteins in myxozoan polar capsule development. Our findings have implications for future comparative and functional studies based upon knowledge of the actual stage of the myxozoan development.

## Materials and methods

### Biological materials and sampling

Northern pikes *E. lucius* (Actinopterygii: Esocidae) were obtained from the fish farm Rybářství Třeboň a.s. (Třeboň, Czech Republic). Fish were euthanised with an overdose of buffered MS-222 (Sigma Aldrich, USA). All animal procedures were performed in accordance with Czech legislation (section 29 of the Protection of Animals Against Cruelty Act No. 246/1992) and approved by the Czech Ministry of Agriculture. Urinary bladders were dissected and screened for the presence of developmental stages of *M. lieberkuehni* using an Olympus BX51 light microscope. The kinetics of myxospore development was checked throughout the year, in 2019 and 2020, while omitting summer months due to the prohibition of pike fishing during this season. For acquisition of stage-specific phases of myxospore development, different *M. lieberkuehni* stages in three technical and three biological replicates were kept in 400 μL TNES urea buffer (10 mM Tris-HCl [pH 8], 125 mM NaCl, 10 mM EDTA, 0.5 % SDS, and 4 M urea) for DNA analysis or in 100 μL RNAlater (Sigma-Aldrich, USA) for RNA extraction. Total genomic DNA was extracted using a standard phenol-chloroform extraction protocol after an overnight incubation with Proteinase K (BioTech a.s., Czech Republic). RNA was extracted using the Total RNA Isolation Kit (Macherey-Nagel, Germany) according to manufacturer’s guidelines. Both the quality and quantity of RNA and DNA were determined by NanoDrop (ThermoFisher, USA). cDNA was synthesized using the Transcriptor High Fidelity cDNA Synthesis Kit (Roche, Switzerland) according to the manufacturer’s instructions.

### PCR validation of minicollagen genes

The transcripts of three minicollagen genes (*ncol-1*, *ncol-3*, *ncol-5*) identified in the de novo transcriptome assembly of *M. lieberkuehni* (Kyslík et al., 2021) were used for primer design (Table S1). Conventional PCR of individual minicollagen genes was carried out by AmpOne HS-Taq premix (GeneAll Biotechnology, South Korea). PCR reactions contained 10 μl of HS-Taq premix, 0.5 μl of each primer (25 pmol), 8 μl of Milli-Q H2O and 1 μl of DNA/cDNA (10 to 150 ng each). Then, the cycling parameters were used as follows: 95 °C 3 min, 35x (95 °C 1 min, 55 °C 1 min, 72 °C 1 min), 72 °C 7 min. PCR products were purified using the Gel/PCR DNA Fragments Extraction Kit (Geneaid Biotech Ltd, Taiwan). The purified products were commercially sequenced (SeqMe s.r.o., Czech Republic). The sequenced amplicons were mapped to individual minicollagen transcripts. Introns, delimited by GT/AG splice sites, were identified by comparison of sequences resulting from DNA and cDNA templates.

### Minicollagen gene expression assays

qPCR gene-specific primers for either minicollagens or reference genes were designed to amplify 80–150 bp regions (Table S1). Primer pairs were designed in the exon-intron gene splice-junction regions with optimal Tm of 58–60 °C and with a GC content between 45–50 %, using the Primer3 (ver. 4.0.1) (Untergasser et al., 2012). The primer specificity was tested using standard PCR prior to RT-qPCR and verified by single amplicon sequencing of the expected size from reverse transcriptase controls (+/-RT). qPCR was performed in technical and biological triplicates using FastStart Universal SYBR™ Green Master Mix (Roche, Switzerland) on the QuantStudio 6 Flex Real-Time PCR System (ThermoFisher, USA). Primer efficiency of each primer pair (Table S1) was evaluated by qPCR on serial cDNA dilutions and was confirmed to be above 95 % and below 110 %. Housekeeping gene selection and validation was performed as previously described (Kosakyan et al., 2019). All gene expression levels were normalised to the levels of β-actin and GAPDH housekeeping genes using the standard curve method and expressed as either fold change (log_10_) or relative gene expression values and visualised in GraphPad (ver. 6).

### Polar capsule extraction

For polar capsule collection, *M. lieberkuehni* myxospores from the urinary tract of *E. lucius* collected from ten fish individuals in April were centrifuged at 1700 RCF for 5 min and incubated in 1 % sodium dodecyl sulfate (SDS) for 60 min while shaking. The following steps were performed, as previously described (Piriatinskiy et al., 2017). Briefly, after SDS treatment samples were washed with 0.1 % Tween 20 and centrifuged at 1000 RCF for 8 min. The pellet was incubated with 1% subtilopeptidase A (Sigma Aldrich, USA) at room temperature (RT) for 60 min followed by two wash steps with 0.05 % Tween 20 each centrifuged at 1000 RCF for 8 min. The final mixture of released polar capsules and spore valves was resuspended in 0.05 % Tween 20 stored at 4 °C for later use.

### Antibodies and Western blot analyses

Custom polyclonal rabbit antibodies against *M. lieberkuehni* Ncol-1 and Ncol-5 were produced against peptides (Ncol-1: AGMPRSIEKRQT; Ncol-5: AGRPGPQGPPGSNSAS) synthetized based on corresponding minicollagen amino acid sequences by Davids Biotechnologie GmbH (Germany). The cross-reactivity of antibodies was verified by their reaction against the synthetized peptides and uninfected fish tissues (pike muscle tissue homogenates). For Western blot analyses, protein samples containing presporogonic or sporogonic *M. lieberkuehni* developmental stages were mixed in 2x Laemmli sample buffer with 10 % 2-β-mercaptoethanol and heated at 97 °C for 5 min before loading on a 12 % (w/v) TGX FastCast SDS/PAGE gel (Bio-Rad, USA). Membranes were blocked in 3 % (w/v) skimmed milk with 0.3 % PBST buffer for 1 hour at RT while shaking. Subsequently, anti-Ncol-1 and anti-Ncol-5 polyclonal antibodies were used at a dilution of 1:200. Membranes were incubated overnight at 4°C and then washed three times with 0.3 % PBST followed by incubation with a horseradish peroxidase (HRP)-conjugated anti-rabbit IgG secondary antibody (Alexa Fluor Plus™ 647 goat anti-rabbit IgG, ThermoFisher, USA; 1:5000) for 1 hour. Membranes were then washed three times with 0.3 % PBST and developed with Clarity Western ECL Substrate (Bio-Rad, USA).

### Predictions of post-translational modifications

The deep-learning predictor MusiteDeep (Wang et al., 2020) was used to predict PTMs. Protein sequences were analysed under default parameters (cut-off: 0.5) with prediction model of all possible PTMs. The outputs were visualized using the same webserver.

### Immunolabelling and imaging

Samples were adhered to Superfrost slides (ThermoFisher, USA) after cytospin for 10 min at 1500 RCF and fixed in 1x paraformaldehyde-containing IC fixation buffer (ThermoFisher, USA) at RT, for 30 min. After rinsing three times in 0.3 % PBST at RT for 5 min, samples were blocked in 5 % (w/v) skimmed milk in 0.3 % PBST at RT for 1 hour and then incubated with anti-Ncol-1 and anti-Ncol-5 primary antibodies (1:100 dilution) at 4°C, overnight, with shaking. Minicollagens were visualised with Alexa Fluor Plus™ 647 goat anti-rabbit IgG (ThermoFisher, USA) at 1:500 dilution. Slides were counterstained with DAPI nuclear stain (Sigma-Aldrich, USA) and mounted in Fluoroshield Mounting Medium (Searle Diagnostic Gurr Products, United Kingdom). Images were captured on an Olympus FLUOVIEW FV1000 confocal laser scanning microscope equipped with FluoView FV31S-SW (Ver.2.6). Optical sections (Z-stacks) were deconvolved using the same software. Negative controls with host red blood cells (RBCs) or with parasite stages without a primary or secondary antibody application were carried out and were consistently negative.

For immunogold electron microscopy (EM), *M. lieberkuehni* developmental stages in urine were fixed by adding an equal volume of the fixative solution (8 % formaldehyde with 0.2 % glutaraldehyde in 0.1 M HEPES) and processed as reported previously (Hartigan et al., 2016). Briefly, samples were washed in 0.01 M glycine in HEPES, embedded in 10 % gelatine at 37 °C, cryoprotected in 2.3 M sucrose at 4°C for 3 days and frozen by plunging into liquid nitrogen. Cryosections were cut with Leica EM FC6 ultramicrotome equipped with Leica UCT cryo-chamber (Leica Microsystems), picked up with 1.15 M sucrose/1 % methylcellulose solution (25 cp, Sigma) and transferred onto Formvar-carbon-coated grids. Sections were blocked for 60 min in the blocking buffer containing 3 % bovine serum albumin (BSA) and 0.05 % Tween 20. Sections were incubated for 30 min with either the custom anti-Ncol-1 (1:40) or anti-Ncol-5 (1:40) antibodies in the blocking buffer, washed and labelled with protein A conjugated to 10 nm gold (CMC Utrecht, 1:25) for 60 min at RT. After washing, sections were post-fixed for 5 min in 1 % glutaraldehyde, washed in dH2O, and then contrasted/embedded using a mixture of 2 % methylcellulose and 3 % aqueous uranyl acetate solution (9:1). Samples were observed using a JEOL TEM 1010.

### Scanning electron microscopy

Extracted polar capsules (20 μl) in 0.05 % PBST were fixed in 3 % glutaraldehyde in 0.1 M cacodylate buffer for 30 min at RT. Samples were then mounted on coverslips coated with poly-L-lysine (0.1 %) for 30 min, then post-fixed with 1 % OsO4, rinsed with distilled water, dehydrated in ascending acetone series (5 min at each step). After critical point drying, coverslips were mounted on stubs, gold-sputtered, and examined using the Apreo 2 SEM (ThermoFisher, USA).

## Acknowledgements

We thank the Czech Science Foundation [19-28399X to ASH/IF, 19-25536Y to AL]. We are grateful to the Ministry of Education, Youth and Sports of the Czech Republic [Czech BioImaging LM2018129 and OP VVV CZ.02.1.01/0.0/0.0/18_046/0016045].

## Author contributions

JK carried out all experiments, analysed data and drafted manuscript. MV performed immunogold-labelling and image interpretation. PB-S advised in gene expression design and data interpretation and edited the manuscript. AL contributed to fish dissection, sample processing, and sample preparation for scanning electron microscopy. ASH contributed to data interpretation and edited the manuscript. IF supervised the study and edited the manuscript. All authors approved the manuscript.

---

**Fig. S1.**
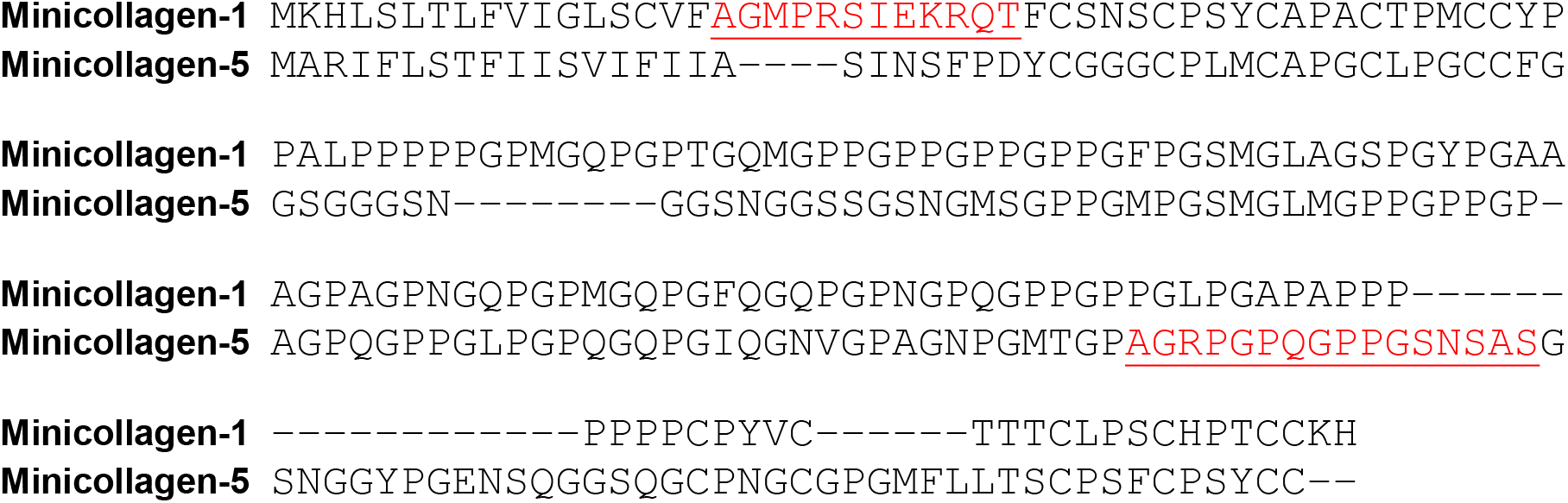
Alignment of *Myxidium lieberkuehni* minicollagen primary amino acid sequences used for design and production of polyclonal antibodies. Peptides selected for de novo peptide synthesis are underlined and highlighted in red.

**Fig. S2.**
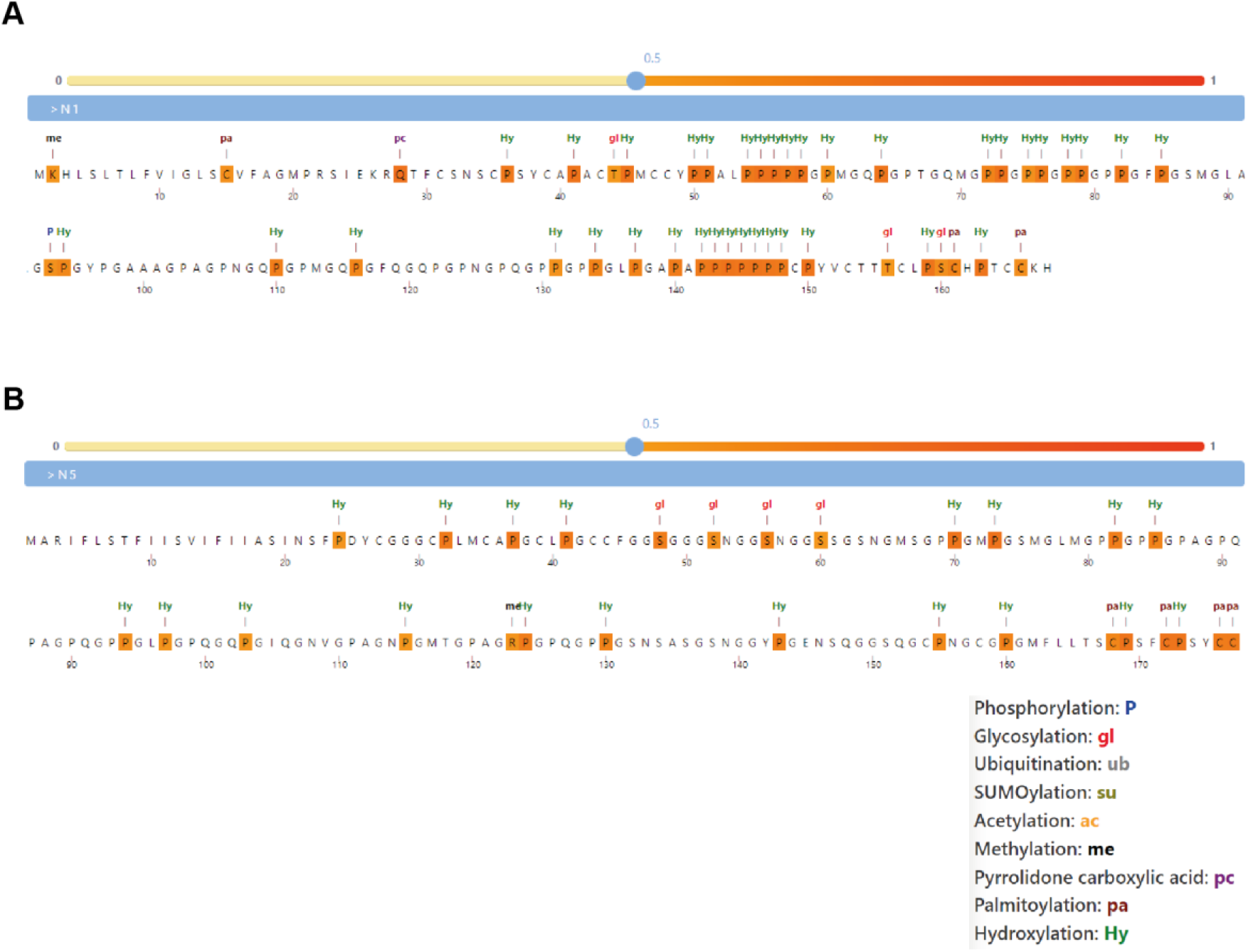
MusiteDeep prediction output of post-translational modifications (PTMs) in *Myxidium lieberkuehni* minicollagens peptides under default cut-off. **(A)** Ncol-1 PTMs prediction output. **(B)** Ncol-5 PTMs prediction output. Individual amino acids with PTMs are highlighted in orange with an indexed abbreviation of predicted PTMs as defined in the legend.

**Fig. S3.**
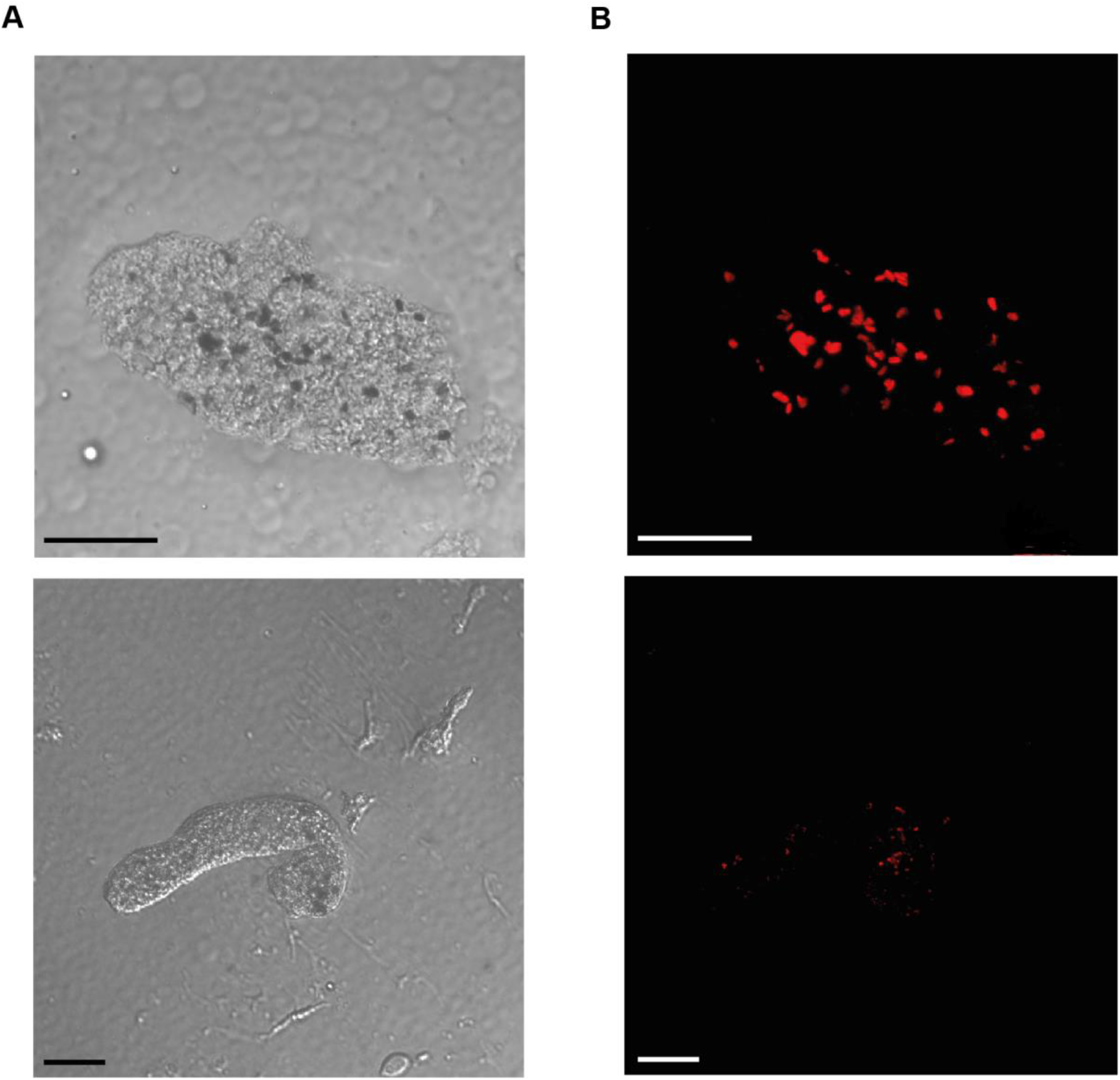
Autofluorescence of crystal-like structures in presporogonic stages (plasmodia) of *Myxidium lieberkuehni*. **(A)** Differential interference contrast images of polysporic plasmodia. **(B)** Epifluorescence images of the same polysporic plasmodia as in (A) with crystal-like structures (red). (Scale bar: 20 μm).

**Table S1.**
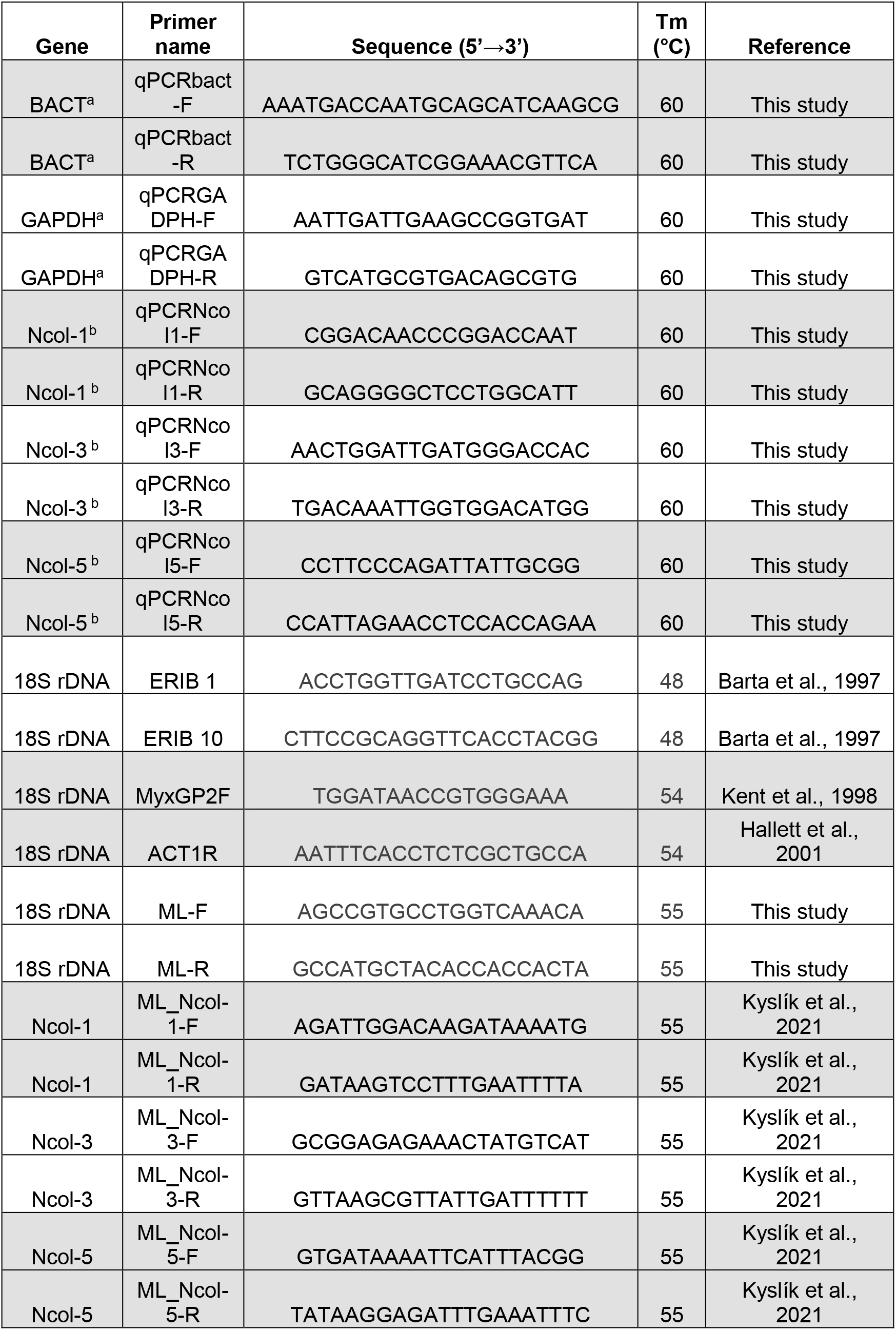
List of primers used for PCR and real time polymerase chain reaction (qPCR) of *M. lieberkuehni* minicollagens and housekeeping genes (HKGs). Primers used as HKGs are indexed by (a), and minicollagen target genes are indexed by (b).

